# Molecular evaluation of *Conogethes punctiferalis* (Guenée, 1854): Species status and intraspecific divergence

**DOI:** 10.1101/2023.10.28.564482

**Authors:** Jin An, Ya-Lin Yao, Ping Gao, Minghua Xiu, Cheng-Min Shi

## Abstract

Species are not only the fundamental units of taxonomy but also the basic units of pest management. Insects of *Conogethes* are important agricultural and forestry pests. However, species boundaries within *Conogethes* often appear obscure. In the present study, we re-evaluated the species status of *Conogethes* by applying three species delimitation approaches based on the mitochondrial DNA sequences, with particular emphasis on the yellow peach moth *C. punctiferalis* (Guenée, 1854). We first optimized species delimitation and inter-species genetic divergence threshold using a DNA barcoding dataset. Our results revealed that several nominal species of *Conogethes* species harbored deeply diverged mitochondrial lineages which were recognized as independent species by the species delimitation methods. The p-distance between the delimited putative species ranged from 0.0159 to 0.1321 with a mean of 0.0841. Then we refined the species status of *C. punctiferalis* using the smallest interspecific distance threshold based on a geographically comprehensive population-scale dataset. This procedure narrowed the species concept of *C. punctiferalis* to a genetically coherent unit. Further investigation of its intraspecific divergence in the geographic context revealed that the refined *C. punctiferalis* was still widely distributed with the same or highly similar mitochondrial haplotypes occurring across South and East Asia.

## 1 Introduction

Species are not only the fundamental unit of taxonomy and biodiversity inventory but also the basic unit of management for those that pose threats to agriculture, forestry, and public health. For pest insects, how species are defined and delimited has profound implications for pest management practices. A basic, but often unstated, assumption of modern pest management is that members of the same target species share similar life cycles, host range, habitat preferences, and other ecological traits that confer similar responses to management tactics. As it has been widely testified that distinct species react differently to control methods, the presence of cryptic pest species (two or more distinct species that are erroneously classified and hidden under one species name) can lead to failure of control efforts for no apparent reason (Walter, 2005). There are many examples where the misidentification of a cryptic species complex as one species has thwarted control efforts. However, most, if not all, agricultural pest insects were initially recognized only from morphological perspectives. Now it is well acknowledged that morphological species do not always present a natural unit that is reproductively isolated from other such units or has a coherent evolutionary history. Cryptic species are common in generalist pest insects, for instance, in the neotropical skipper butterfly *Astraptes fulgerator* (Hebert et al., 2004), and the whitefly *Bemisia tabaci* (Vyskočilová et al., 2018). The consequence of cryptic species on pest management is particularly profound for species that have developed insecticide resistance which appears common in agroecosystems (Horowitz et al., 2020).

The yellow peach moth *Conogethes punctiferalis* (Guenée, 1854), belonging to the specious family Crambidae (Insecta: Lepidoptera), is an important agricultural pest. It was first described as *Astura punctiferalis*, then revised as *Dichocrocis punctiferalis* before getting its current name. It is widespread in subtropical and tropical Asia and Oceania, and of particular concern in agriculture in China, India, Indonesia, Korea, Japan, Malaysia, New Guinea, Vietnam and Australia (EPPO, 2023). This species is highly polyphagous feeding on more than 100 host plants from >30 different families, many of which are economically important crops and fruits (Lu et al., 2010; Rojas-Sandoval, 2022). Larvae of *C. punctiferalis* have long been considered a serious pest of peach, apple, pear, chestnut and other fruits across its range, and recently emerged as a devastating pest of maize and sunflower in north China (Wang et al., 2006). The wide geographic distribution combined with extreme polyphagy implies cryptic divergence within the species.

Fueled by the increasing availability of DNA sequences, the discoveries of cryptic species have increased exponentially over the past three decades (Bickford et al., 2007). Indeed, two ecotypes of *C. punctiferalis* with different host plant preference observed in Japan was recognized as two distinct species (Honda & Mitsuhashi, 1989). An oligophagous Pinaceae-feeding type (PFT) mainly feeding on pines was recognized as *C. pinicolalis*; a still polyphagous fruit-feeding type (FFT) was re-described as *C. punctiferalis* (Inoue & Yamanaka, 2006). The validity of the recognized species was recently supported by divergence in morphology, ecology and genetics in terms of both mitochondrial DNA and nuclear DNA (Jeong et al., 2021). Substantial genetic divergences were detected between the two species with an average of 5.46% for the mitochondrial COI genes and 2.10% for the nuclear EF1α sequences (Jeong et al., 2021). Both of these two species, referred to here as *C. punctiferalis s. str*. and *C. pinicolalis*, were identified in China based on mitochondrial DNA sequences (Wang et al., 2014). Similarly, *C. punctiferalis* in India feeding on castor and cardamom represented two different lineages (Chakravarthy et al., 1991). The cardamom population was lately recognized as a distinct species *C. sahyadriensis* which differed from *C. punctiferalis* by genetic distances of no less than 6.00% (Shashank et al., 2018). Thus, it is clear that *C. punctiferalis* represents a species complex including multiple species of very similar morphology, variable ecotypes, and overlapping host ranges (Armstrong, 2010).

Although several cryptic species with restricted host plants and narrow geographic range have been recognized in the traditional morphological species *C. punctiferalis*, its remaining members (*C. punctiferalis s. str*.) still appear extremely host-rich and widespread. To better inform the management of this pest insect in the future, a global re-evaluation of its species status and intraspecific genetic variation is warranted. Here we performed such a molecular re-evaluation based on mitochondrial DNA sequence data. We first delimitated species by applying three statistical species delimitation methods and optimized species recognition threshold. Using the threshold obtained, we re-evaluated the species status and intraspecific genetic variation of *C. punctiferalis s. str*. based on a global dataset.

## 2 Materials and methods

### 2.1 Data curation

Two DNA datasets of the mitochondrial COI gene (mtCOI) were compiled in the present study. The first dataset, referred to as ConCOI, was used to examine the performance of species delimitation methods and thereby optimize the threshold of genetic distance for species recognition. The dataset included mtCOI sequences for all species of *Conogethes* currently recognized. We derived the ConCOI dataset by querying the GenBank through BLASTn with mtCOI sequences of the putative species recognized by the taxonomic treatment of Shashank et al. (2018). We filtered the hits from each of blast search with identity of 95%∼100% and coverage of 98%∼100%. Sequences were aligned using MUSCLE (Edgar, 2004). The aligned ConCOI dataset included 297 mtCOI sequences of *Conogethes* species and one sequence (HQ952111) for *Oxycanus niphadias* as an outgroup. After trimming, the 297 mtCOI sequences of *Conogethes* species harbored 110 unique sequences (haplotypes) of 566-bp in length.

The second dataset, referred to as CpunCOI, was used to re-evaluation species status and intraspecific genetic variation. This dataset was compiled by downloading all available mtCOI with taxon label of *Conogethes punctiferalis* from GenBank and BOLD databases. After filtering using the within-species genetic distance of *C. punctiferalis* inferred from the ConCOI dataset (see below), the CpunCOI dataset included 420 mtCOI sequences of 233 bp long after trimming to the same length. We preferred to include more individuals over selecting longer sequences as it has been demonstrated that there were no significant differences in performance for species-level identification between full-length and mini-barcodes as long as they are > 200 bp in length (Yeo et al., 2020).

### 2.2 Tests of species limits and optimization of genetic threshold

We tested species limits in *Conogethes* using the ConCOI haplotype dataset by employing both the genetic clustering method and the phylogenetic tree-based approaches. Our contention is that robust species delimitations will be consistently supported by signals from multiple species delimitation approaches. For the genetic clustering method, we used the assembling species by automatic partitioning (ASAP) method which employs pairwise genetic distances for hierarchical clustering (Puillandre et al., 2021). The genetic distances were computed using K80 (a.k.a. K2P) substitution model with the transition vs. transversion ratio (ts/tv = 3.74) being estimated from the data using the maximum likelihood method. The best species partition was identified according to the ASAP score. Following the recommendation of Puillandre et al. (2021), we considered the partitions with the top two best ASAP scores.

Two tree-bases methods were applied. First, potential species were delimited using the generalized mixed Yule Coalescent (GMYC) method (Fujisawa & Barraclough, 2013) based on the maximum clade credibility (MCC) tree inferred using BEAST v.1,10.4 (Suchard et al., 2018). We only used the single threshold model because we expected the result to be transferable among datasets. BEAST was run with TN93+ G4 substitution model and strict molecular model with a Yule process as speciation prior. Markov chain Monte Carlo (MCMC) searches were run for 2 × 10^7^ generations with trees and parameters being sampled every 2000 generations. We used Tracer 1.7 (Rambaut et al. 2018) to check the stationarity and the convergence of the MCMC chains. MCC tree was summarized from the sampled trees using TreeAnnotator 1.10 (Suchard et al. 2018), with the first 10% of samples being discarded as burn-in. Second, species were delimited using the Poisson tree process (PTP) model (Zhang et al., 2013) based on the maximum likelihood tree inferred using IQ-TREE v. 2.3 (Nguyen et al., 2015) and applying a single threshold. We used both the maximum likelihood and Bayesian implementation of the PTP model (mPTP and bPTP, respectively). The analysis was run for 100000 generations with a thinning parameter of 100 and burn-in of 0.1.

Pairwise genetic distances (*p*-distance and K2P-distance) among afore-delimited species were calculated using MEGA 11(Tamura et al., 2021). The smallest interspecific distance was used as a putative genetic threshold for further demarcation of species boundary of *Conogethes* species. It has been reported that the use of the mean distance exaggerates the size of interspecific divergences and leads to misidentification (Meier et al., 2008)

### 2.3 Analyses of intraspecific genetic variations for *C. punctiferalis s. str*

For the clade/cluster or delimited species corresponding to *C. punctiferalis* (sensu Shashank et al. 2018), the pairwise genetic distances (*p*-distance) among mtCOI haplotypes were calculated using MEGA 11(Tamura et al., 2021). To further evaluate intraspecific genetic variation, we used these haplotypes as queries and recursively blasted the GenBank with the smallest interspecific distance found among the species delimited in the present study as a filtering threshold. After removing repeated sequences, aligning with above haplotypes and trimming to the same length, these sequences made up of the CpunCOI dataset. We determined the unique haplotypes and calculated population genetic summary statistics using DnaSP version 6.12.03 (Rozas et al., 2017). Median-joining networks (Bandelt et al., 1999) were constructed using NETWORK 10.2.0.0 (https://www.fluxus-engineering.com/) to illustrate intraspecies relationships, and geographic distribution of unique haplotypes.

## 3 Results

### 3.1 Delimitation of *Conogethes* species

The ConCOI dataset included 297 mtCOI sequences (566 bp) which were collapsed into 110 unique haplotypes. From this dataset, the number of species delimitation by three different methods ranged from 15 to 23 (Figure 1). The best partition found by ASAP clustered the 110 haplotypes of *Conogethes* into 19 subsets (species) with an ASAP score of 4.00 and a distance threshold of 0.0143 (ASAP-1 in Figure 1). The second-best partition (ASAP score = 4.50) clustered the *Conogethes* haplotypes into 14 subsets with a distance threshold of 0.0343 (ASAP-2 in Figure 1). The maximum likelihood partition found by the maximum likelihood search of the PTP model (mPTP) delimited 20 species. The most supported partition found by Bayesian PTP search (bPTP) delimited 21 species. One species (ANIC631-06, HQ953112) delimited by mPTP was split into two independent species. We note that these two sequences were poorly differentiated in the Bayesian MCC tree (Figure 1). Applying the single threshold, GMYC delimited 23 species (confidence interval: 18∼27). The species delimitation model was significantly better than the null model (likelihood ratio: 26.60, result of LR test: 1.67e-06). The consensus species delimitation among the three methods recognized 19 species (Figure 1). All of these species formed strongly supported monophyletic clades. Except for *C. pinicolalis*, all other 18 species included identical or highly similar mtCOI sequences to those used by recent taxonomic treatment by Shashank et al. (2018), so we named each species following Shashank et al. (2018) to make it easy for cross-references.

**Figure 1.**
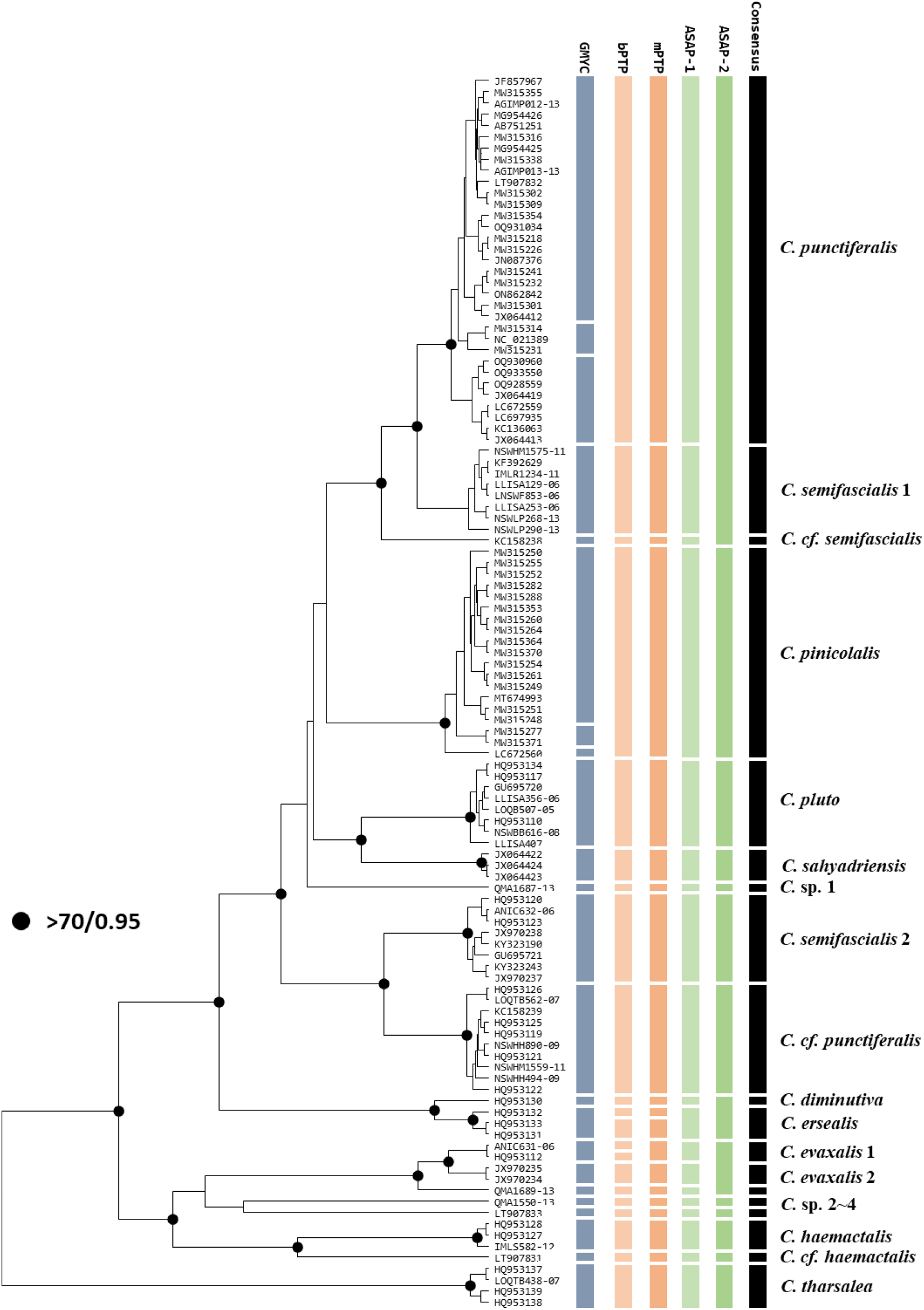
Species delimitation and phylogenetic relationship of *Conogethes* mitochondrial haplotypes. The ultrametric phylogenetic tree was inferred using BEAST with a strict molecular clock model and a speciation prior of Yule process. Strongly supported nodes with posteriors ≥ 0.95 and maximum likelihood bootstrapping support values ≥ 70 were indicated by black dots. The bars on the right depict the putative species delimited by different methods. Species names identified following Shashank et al. (2018).

The genetic distances calculated from mtCOI sequences of the 19 recognized consensus species are summarized in Table 1. The interspecific p-distances ranged from 0.0159 to 0.1321, with a mean of 0.0841. The corresponding K2P distances were 0.0161 to 0.1455, and 0.900, respectively. The maximum within species genetic distances were 0.0066 and 0.0067 for p-distances and K2P distances, respectively.

**Table 1.** Estimates of genetic divergence between the delimited species of *Conogethes*. The p-distances were shown above the diagonal and the K2P distances were shown the diagonal. The numbers on the diagonal represented the within species K2P distances.

| Species | Cpun | Csem1 | Ccfsem | Cpin | Csah | Ccfpun | Csp1 | Csem2 | Cplu | Ccfhae | Cers | Ceva2 | Csp2 | Cdim | Chae | Csp4 | Ceva1 | Ctha | Csp3 |
| --- | --- | --- | --- | --- | --- | --- | --- | --- | --- | --- | --- | --- | --- | --- | --- | --- | --- | --- | --- |
| **Cpun** | **0.0066** | 0.0230 | 0.0365 | 0.0500 | 0.0518 | 0.0519 | 0.0549 | 0.0561 | 0.0591 | 0.0791 | 0.0815 | 0.0850 | 0.0859 | 0.0856 | 0.0894 | 0.0887 | 0.0931 | 0.0974 | 0.1078 |
| **Csem1** | 0.0235 | **0.0024** | 0.0318 | 0.0503 | 0.0496 | 0.0409 | 0.0526 | 0.0480 | 0.0570 | 0.0773 | 0.0699 | 0.0813 | 0.0808 | 0.0775 | 0.0855 | 0.0797 | 0.0866 | 0.0965 | 0.1005 |
| **Ccfsem** | 0.0375 | 0.0326 | **n.c.** | 0.0547 | 0.0624 | 0.0481 | 0.0601 | 0.0590 | 0.0548 | 0.0901 | 0.0789 | 0.0936 | 0.0972 | 0.0883 | 0.0954 | 0.0954 | 0.0919 | 0.1056 | 0.1113 |
| **Cpin** | 0.0522 | 0.0526 | 0.0572 | **0.0060** | 0.0615 | 0.0636 | 0.0592 | 0.0603 | 0.0561 | 0.0810 | 0.0664 | 0.0801 | 0.0849 | 0.0751 | 0.0863 | 0.0951 | 0.0872 | 0.1100 | 0.1025 |
| **Csah** | 0.0544 | 0.0521 | 0.0661 | 0.0651 | **0.0012** | 0.0691 | 0.0607 | 0.0710 | 0.0527 | 0.0942 | 0.0760 | 0.0989 | 0.0954 | 0.0819 | 0.0901 | 0.1013 | 0.1042 | 0.0991 | 0.1119 |
| **Ccfpun** | 0.0545 | 0.0424 | 0.0500 | 0.0671 | 0.0738 | **0.0034** | 0.0641 | 0.0408 | 0.0593 | 0.0852 | 0.0690 | 0.1019 | 0.0966 | 0.0834 | 0.0908 | 0.0905 | 0.0970 | 0.1042 | 0.1108 |
| **Csp1** | 0.0575 | 0.0550 | 0.0631 | 0.0621 | 0.0640 | 0.0677 | **n.c.** | 0.0643 | 0.0526 | 0.0954 | 0.0736 | 0.1025 | 0.0972 | 0.0813 | 0.0872 | 0.0954 | 0.1042 | 0.1038 | 0.1095 |
| **Csem2** | 0.0589 | 0.0500 | 0.0620 | 0.0633 | 0.0757 | 0.0424 | 0.0677 | **0.0044** | 0.0654 | 0.0905 | 0.0862 | 0.1011 | 0.1003 | 0.0939 | 0.0864 | 0.0952 | 0.1029 | 0.1080 | 0.1100 |
| **Cplu** | 0.0625 | 0.0603 | 0.0575 | 0.0591 | 0.0557 | 0.0626 | 0.0550 | 0.0693 | **0.0025** | 0.1014 | 0.0674 | 0.1009 | 0.1009 | 0.0800 | 0.0917 | 0.0972 | 0.0961 | 0.1127 | 0.1133 |
| **Ccfhae** | 0.0837 | 0.0816 | 0.0961 | 0.0858 | 0.1009 | 0.0905 | 0.1022 | 0.0966 | 0.1093 | **n.c.** | 0.0907 | 0.0813 | 0.0777 | 0.1007 | 0.0595 | 0.0883 | 0.0830 | 0.1038 | 0.1007 |
| **Cers** | 0.0871 | 0.0739 | 0.0842 | 0.0701 | 0.0809 | 0.0730 | 0.0780 | 0.0927 | 0.0712 | 0.0969 | **0.0035** | 0.0995 | 0.1013 | 0.0230 | 0.0801 | 0.0978 | 0.0942 | 0.1104 | 0.1037 |
| **Ceva2** | 0.0902 | 0.0861 | 0.1002 | 0.0847 | 0.1064 | 0.1099 | 0.1105 | 0.1089 | 0.1088 | 0.0861 | 0.1074 | **0.0000** | 0.0247 | 0.1042 | 0.0878 | 0.0813 | 0.0159 | 0.1140 | 0.0989 |
| **Csp2** | 0.0913 | 0.0856 | 0.1043 | 0.0903 | 0.1023 | 0.1037 | 0.1043 | 0.1078 | 0.1088 | 0.0821 | 0.1095 | 0.0253 | **n.c.** | 0.1060 | 0.0895 | 0.0901 | 0.0318 | 0.1184 | 0.1007 |
| **Cdim** | 0.0919 | 0.0827 | 0.0952 | 0.0799 | 0.0877 | 0.0896 | 0.0868 | 0.1018 | 0.0855 | 0.1086 | 0.0235 | 0.1131 | 0.1152 | **n.c.** | 0.0836 | 0.1007 | 0.1095 | 0.1184 | 0.1095 |
| **Chae** | 0.0952 | 0.0909 | 0.1022 | 0.0917 | 0.0961 | 0.0970 | 0.0927 | 0.0919 | 0.0979 | 0.0621 | 0.0849 | 0.0934 | 0.0955 | 0.0889 | **0.0024** | 0.0819 | 0.0919 | 0.1092 | 0.0948 |
| **Csp4** | 0.0945 | 0.0843 | 0.1023 | 0.1020 | 0.1093 | 0.0966 | 0.1023 | 0.1020 | 0.1044 | 0.0942 | 0.1053 | 0.0861 | 0.0962 | 0.1087 | 0.0869 | **n.c.** | 0.0866 | 0.1113 | 0.0989 |
| **Ceva1** | 0.0996 | 0.0921 | 0.0981 | 0.0929 | 0.1127 | 0.1041 | 0.1126 | 0.1109 | 0.1031 | 0.0881 | 0.1012 | 0.0161 | 0.0327 | 0.1195 | 0.0982 | 0.0922 | **0.0000** | 0.1170 | 0.1042 |
| **Ctha** | 0.1045 | 0.1034 | 0.1139 | 0.1191 | 0.1065 | 0.1124 | 0.1120 | 0.1168 | 0.1226 | 0.1118 | 0.1199 | 0.1238 | 0.1290 | 0.1296 | 0.1182 | 0.1207 | 0.1274 | **0.0041** | 0.1321 |
| **Csp3** | 0.1166 | 0.1080 | 0.1207 | 0.1104 | 0.1217 | 0.1202 | 0.1186 | 0.1192 | 0.1234 | 0.1083 | 0.1118 | 0.1063 | 0.1084 | 0.1188 | 0.1017 | 0.1065 | 0.1126 | 0.1455 | **n.c.** |

### 3.2 Intraspecific genetic divergence of *Conogethes punctiferalis*

The p-distances between 33 mtCOI haplotypes (566 bp) of *C. punctiferalis* from ConCOI dataset ranged from 0.0018 to 0.0159 with an average distance of 0.0066 (Table 2). To further evaluate within species genetic divergence of *C. punctiferalis*, we filtered the CpunCOI dataset with the minimum inter-species genetic distance found among the delimited species (0.0159, *C. evaxalis* 1 vs. *C. evaxalis* 2; Table 1). The final CpunCOI dataset included 420 sequences of 233 bp long. These 420 sequences harbored 39 haplotypes defined by 30 variable sites. The haplotype diversity and nucleotide diversity were 0.6100 and 0.0053, respectively. The p-distances between the 39 short haplotypes (233 bp) of *C. punctiferalis* ranged from 0.0043 to 0.0343 with an average of 0.0126 (Table 2).

**Table 2.** Estimates of genetic divergence between mtCOI haplotypes of *C. punctiferalis*. n: sample size; H: number of haplotypes.

| Datasets | Length | n | H | p-distance (mean [min., max.]) | K2P-distance (mean [min., max.]) |
| --- | --- | --- | --- | --- | --- |
| ConCOI | 566 bp | 147 | 33 | 0.0066 [0.0018, 0.0159] | 0.0067 [0.0018, 0.0161] |
| CpuCOI | 233 bp | 420 | 39 | 0.0126 [0.0043, 0.0343] | 0.0128 [0.0043, 0.0352] |

The relationships among haplotypes are shown by the median-joining network in Figure 2. The haplotype frequencies range from 1 to 257 (H02). Twenty-two haplotypes are singletons that appeared only once in the dataset. The majority of haplotypes were connected to the dominant haplotype H02 by one mutation step. The second most frequent haplotype (H14; frequency, 44) was connected to H02 by three mutation steps and two intermediate haplotypes. The 39 haplotypes were found in six countries (China, Japan, South Korea, India, Pakistan and Malaysia). Ten haplotypes were shared at least by two counties. Haplotype H02 was the most widely spread one, occurring in all six countries. Haplotype H14 was found in South Korea, China, India, Pakistan and Japan. Haplotypes H04, H08, H11and H15 were shared by East Asian countries (China, Japan and South Korea). Haplotype H03 and H18 were shared between East Asia (China and/or Japan and South Korea) and South Asia (India). In summary, we found extremely wide geographic distributions of the same or highly similar mtCOI haplotypes of *C. punctiferalis* across South and East Asia.

**Figure 2.**
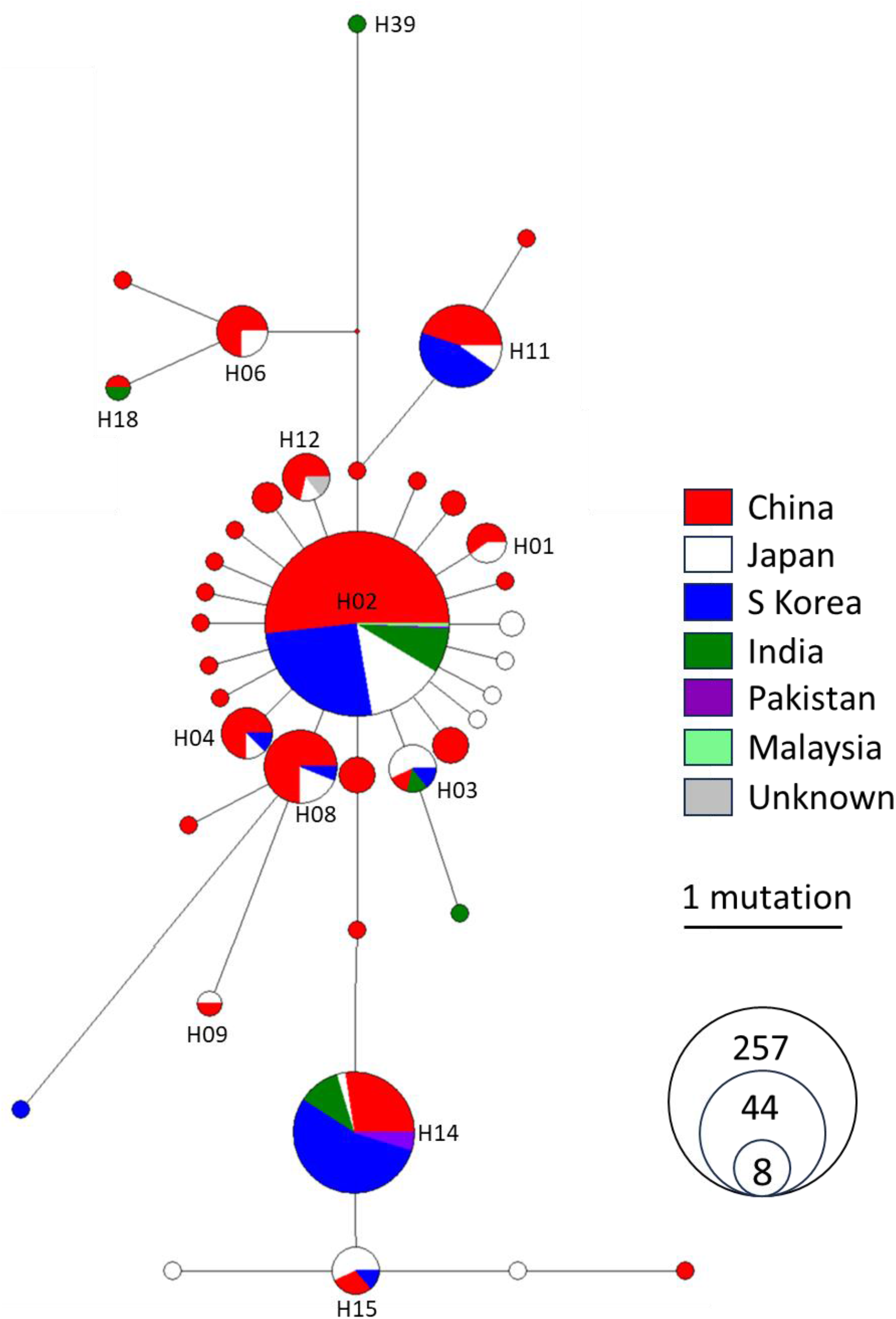
Median-joining network of mitochondrial haplotypes of refined *Conogethes punctiferalis*. Each node represents a unique haplotype identified from 420 mtCOI sequences. Node size is proportional to the frequency of relevant haplotypes, and branch length is proportional to the mutations that occurred between haplotypes. Color depicts the geographic origins of the harboring mtCOI sequences.

## 4 Discussion

### 4.1 Multiple divergent mitochondrial lineages challenge species recognition of *Conogethes*

Species are often treated as the basic unit in the management of pest insects and cryptic species present a challenge to pest management practice (Walter, 2005). It has been long suggested that some members of *Conogethes* represent species complexes including multiple morphologically similar species (Armstrong, 2010). However, species boundaries for most members of *Conogethes* remain to be systematically evaluated using a consistent standard. Here, we bridged this knowledge gap by performing molecular species delimitation using a comprehensive dataset that included all available mtCOI sequences of *Conogethes* species. Species delimitation using three different approaches (5 methods) consistently revealed that several nominal species of *Conogethes* included multiple highly diverged mitochondrial lineages which warrant independent species (Figure 1). For instance, nominal species *C. punctiferalis* was divided into two distantly related monophyletic clades (*C. punctiferalis* vs. *C*.*cf. punctiferalis*) which differed by 5.91% in mtCOI sequences (Table 1). Similarly, the nominal species *C. semifascialis* was divided into three deeply diverged lineages (*C. semifascialis* 1, *C. semifascialis* 2 and *C. cf. semifascialis*). Besides, several deeply diverged mtCOI lineages (*C*. sp. 1∼4), potentially representing distinct species, await to be formally described. Unexpectedly, we found *C. punctiferalis* was more closely related to *C. semifascialis* 1 than to any species that were recently recognized in the *C. punctiferalis* species complex, i.e. *C. pinicolalis* and *C. sahyadriensis* (Figure 1). These findings together signified that DNA-based identification of *Conogethes* species is essential for precision pest management practice for this group.

Considering the consensus species delimitation reported here and species recognition by Shashank et al. (2018) as valid, the minimum inter-species genetic difference was as low as 1.59%, although the overall average of pairwise between-species genetic difference was more than 8% (Table 1). This observation reinforced the earlier finding that use of mean instead of the smallest interspecific distance exaggerates the threshold of the barcoding gap and leads to the over-lumping of species (Meier et al., 2008). We thus suggest the use of the smallest inter-species genetic distance (1.59%) as a putative threshold for molecular recognition of *Conogethes* pest species in management practice.

### 4.2 Redefine species concept of *Conogethes punctiferalis*

The two datasets (ConCOI and CpunCOI) compiled in the present study allowed us to refine the species boundary of *C. punctiferalis* from genetic and geographic perspectives. From a phylogenetic point of view, we redefined *C. punctiferalis* as a strongly supported monophyletic mitochondrial lineage which was most closely related to *C. semifascialis* 1 (Figure 1). The genetic distance between *C. punctiferalis* and its most close relative was 2.30% (p-distance, and 2.35% for K2P distance). Only the GMYC method delimited two sub-lineages within *C. punctiferalis* as independent species (Figure 1). However, these sub-lineages were not well supported in the phylogenetic analyses. Thus, we treated them as intraspecific variations. The species concept of *C. punctiferalis* redefined here was consistent with Shashank et al. (2018).

From a geographic perspective, the refined *C. punctiferalis* is still widely spread. It occurred in China, Japan, South Korea, India, Pakistan, and Malaysia, straddling the temperate, subtropical and tropical zones. The cross-regional sharing of the same mitochondrial haplotypes suggests that this species may have high dispersal capacity and that long-distance dispersal might have been rampant. Such possibility was also supported by the detection of *C. punctiferalis* in the autumn migrating insect troops across northern China by radar observatories (Feng et al., 2003). Therefore, we think *C. punctiferalis* we redefined here represents a coherent genetic unit which may withstand tests from other aspects.

### 4.3 Limitations and perspectives

We should note that we have delimitated species *Conogethes* species, and *C. punctiferalis* in particular, only from a maternal (mitochondrial) perspective. Now, it is increasingly recognized that mitochondrial DNA records the incomplete history of species (Ballard & Whitlock, 2004), and in some cases, mitochondrial DNA can mislead species recognition (Rubinoff & Holland, 2005). Despite the aforementioned and other obvious drawbacks of using mitochondrial DNA alone for species delimitation, the success of using mitochondrial DNA in distinguishing species from a range of taxa and revealing cryptic species is remarkable (Dasmahapatra & Mallet, 2006). Our results therefore can serve as a working hypothesis for further tests. Future efforts should be made to delimitate *Conogethes* species using nuclear DNA data and more robust methods that consider both the speciation process between species and the coalescence process within species (Rannala & Yang 2020). As the advantages and limitations of DNA-based species identification become apparent in pest management practice, it is clear that taxonomic approaches integrating DNA data, morphological characteristics, ecological features, and damage symptoms will achieve optimal efficiency in species identification.

